# Dissecting FcγR Regulation Through a Multivalent Binding Model

**DOI:** 10.1101/142273

**Authors:** Ryan A. Robinett, Ning Guan, Anja Lux, Markus Biburger, Falk Nimmerjahn, Aaron S Meyer

## Abstract

Many immune receptors transduce activation across the plasma membrane through their clustering. With Fcγ receptors, this clustering is driven by binding to antibodies of differing affinities that are in turn bound to multivalent antigen. As a consequence of this activation mechanism, accounting for and rationally manipulating IgG effector function is complicated by, among other factors, differing affinities between FcγR species and changes in the valency of antigen binding. In this study, we show that a model of multivalent receptor-ligand binding can effectively account for the contribution of IgG-FcγR affinity and immune complex valency. This model in turn enables us to make specific predictions about the effect of immune complexes of defined composition. In total, these results enable both rational immune complex design for a desired IgG effector function and the deconvolution of effector function by immune complexes.

**Summary points:**

- Avidity most prominently modulates low-affinity FcγR-immune complex binding
- A multivalent binding model can quantitatively predict FcγR-immune complex binding
- Immune complex avidity has an outsized contribution to FcγR multimerization as compared to binding
- A binding model deconvoles and predicts the influence of interventions modulating *in vivo* FcγR-driven effector function

## Introduction

Antibodies are critical and central regulators of the immune response. Antibodies of the IgG isotype interact with FcγR receptors expressed widely on innate immune effector cells. IgGs transduce effector function through multiple cell types—including macrophages, monocytes, neutrophils, and NK cells—and through multiple processes, including promoting antibody-dependent cell-mediated cytotoxicity (ADCC), antigen presentation, and cytokine response. IgG immunotherapies, operating through regulating effector cell function, have been used in the treatment of both cancer and autoimmune diseases. In cancer treatment, IgG therapies can show a synergistic effect when used in combination with checkpoint or cytokine-mediated immunotherapies^1,2^. These biologic agents are particularly versatile therapeutic agents on account of their immunotherapeutic effects and their ability to operate directly through antigen binding and opsonization.

The ability to quantitatively predict FcγR-IgG function would aid the understanding and treatment of cancer, autoimmune diseases, and infectious diseases. Efforts to engineer IgG treatments with improved effector response have included designing Fc variants with biased FcγR binding, deglycosylating Fc domains (with the effect of modulating FcγR binding), and utilizing alternative IgG subclasses with distinct binding profiles^3,4^. In cases where antigen and antibody are exogenously provided, avidity and binding affinities may be manipulated coordinately in a controlled manner^5^. With a better understanding of the underlying regulation, endogenous humoral responses might similarly be modulated through adjuvant engineering^6^.

Previous efforts have sought to improve our understanding of IgG-mediated effector function. These include efforts to carefully quantify the individual, monovalent FcγR-IgG interaction affinities^7–9^. Others have characterized the effects of IgG glycosylation (which serves to modulate FcγR affinity) and immune complex (IC) avidity on the binding of IgG-antigen complexes^5,10^. Genetic models have made it possible to remove certain FcγRs and examine the consequent effect on IgG treatment, including in the treatment of various cancers^11–13^. By comparing antibodies of matched variable region but differing Fc domains, one can evaluate the influence of effector function, though with necessarily pleiotropic effects on binding to each FcγR class^12,13^.

Models of multivalent ligand binding to monovalent receptors have been successfully employed to study the function of other immune receptors with corresponding binding models^14–16^. For example, a two-component binding model can capture the effect of T cell receptor activation or FcॉRI binding^17,18^. However, unlike these cases, distinct members of the FcγR family can be simultaneously expressed within most cells. Additionally, the multiple FcγRs present, with activating and inhibitory roles, ensure that any manipulation of IC composition will necessarily have multivariate effects. Thus, while the underlying theoretical models of multivalent binding are long-standing, FcγR-IgG interactions are especially suited for developments in inference approaches to rigorously link these models to experimental observations^19–21^.

In this study, we have employed a model of multivalent IC binding to FcγRs and show that it can capture experimentally measured binding at differing valencies. Applying this model, we show it can quantitatively predict effector response to diverse interventions in a forward manner and can deconvolve the causal factors of response in a reverse fashion. More broadly, these results demonstrate the abilities of both a unified binding model and computational inference techniques to link theory and experimental observation.

## Results

### IgG-FcγR binding varies with affinity and avidity

Building upon earlier work in which IC avidity was shown to alter human FcγR-IgG binding, we wished to examine the influence of IC avidity and IgG composition on the binding and activation of different human FcγRs (hFcγRs)^10^. We assessed the binding of ICs presenting one of four human IgG (hIgG) subclasses to cells expressing one of six hFcγR subclasses at a single IC concentration. To do so, we utilized a panel of CHO cell lines, each stably expressing a single hFcγR species, and two populations of IC, with averaged avidities of four and 26, assembled by covalently attaching 2,4,6-trinitrophenol (TNP) to bovine serum albumin (BSA; see Methods). Anti-TNP antibodies of differing hIgG subclass were bound to the BSA complexes before treatment. The measured binding and variation in binding with avidity recapitulated that measured before, with variation as a function of hIgG subclass, hFcγR, and avidity^10^ (Fig. 1A).

**Figure 1:**
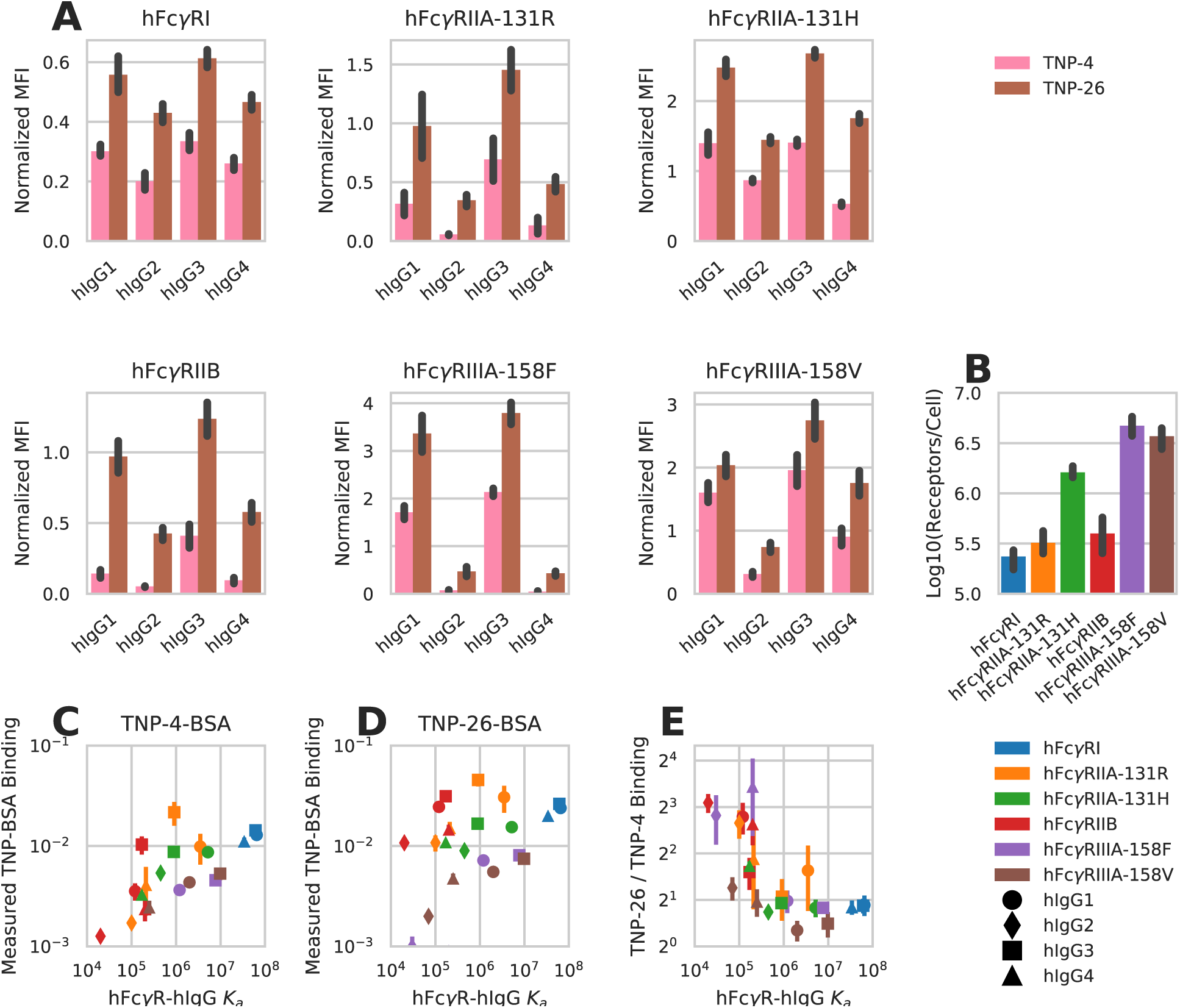
Human FcγR binding changes with FcγR-IgG pair and avidity. A) Qantifcation of hIgG subclass TNP-4-BSA and TNP-26-BSA IC binding to CHO cells expressing the indicated hFcγRs (N = 4). Background binding of the ICs to CHO cells expressing no hFcγR was subtracted from the mean fuorescence intensity (MFI) obtained from binding to CHO cells expressing individual hFcγRs. Each IC binding measurement was furter normalized by dividing by the average of all the points within that replicate. B) Receptor expression quantifcation for each CHO cell line expressing a single hFcγR subclass. C) Measured TNP-4-BSA-IC binding, normalized to the receptor expression within each CHO cell line, as a function of the measured hFcγR-hIgG subclass affinity^7^. D) Measured TNP-26-BSA-IC binding, normalized to the receptor expression within each CHO cell line, as a function of the measured hFcγR-hIgG subclass affinity^7^. E) Fold increase in TNP-26-BSA binding over TNP-4-BSA binding as a function of the measured hFcγR-hIgG subclass affinity. All error bars are standard error of biological replicates (N <= 4). Derived quantities use error propogated from each value.

We quantitatively measured receptor abundance for each hFcγR-expressing cell line to account for this potential source of variation in binding. This revealed twenty-fold variation in the amount of each hFcγR expressed (Fig. 1B). To interpret these measurements, we normalized the amount of binding measured to the amount of hFcγR expressed and plotted each measurement against the measured affinity of the individual hFcγR-hIgG monovalent interaction. We observed a strong correlation between the affinity of the relevant hFcγR-hIgG pair and measured binding, along with a shift toward more binding with increased avidity (Fig. 1C-D). By comparing each TNP-26-BSA and TNP-4-BSA measurement, we observed that the avidity-dependent change in binding varied with the relevant affinity of the hFcγR-hIgG monovalent interaction (Fig. 1E). Lower afn-ity interactions were more avidity-dependent, and the measured binding for each IC with low monovalent affinity was too high to be the result of monovalent binding alone. These observations indicated to us that affinity, avidity, and receptor expression were all critical to interpreting IC binding.

### A multivalent interaction model accounts for variation in FcγR-IgG binding

To interpret the complicated variation in binding we observed with each hFcγR-hIgG pair (i.e. affinity), receptor expression, and IC avidity, we employed an equilibrium model of multi-valent ligand/monovalent receptor binding^17^. Within the model, an initial binding event occurs with the kinetics of the monovalent interaction (Fig. 2A). Subsequent binding events with the same IC occur with a partition coefficient (crosslinking parameter) *K*_*x*_ (see Methods). Values of *K*_*x*_ much greater than 1 promote highly multivalent interactions, while *K*_*x*_ values much less than 1 result in predominantly monovalent binding.

**Figure 2:**
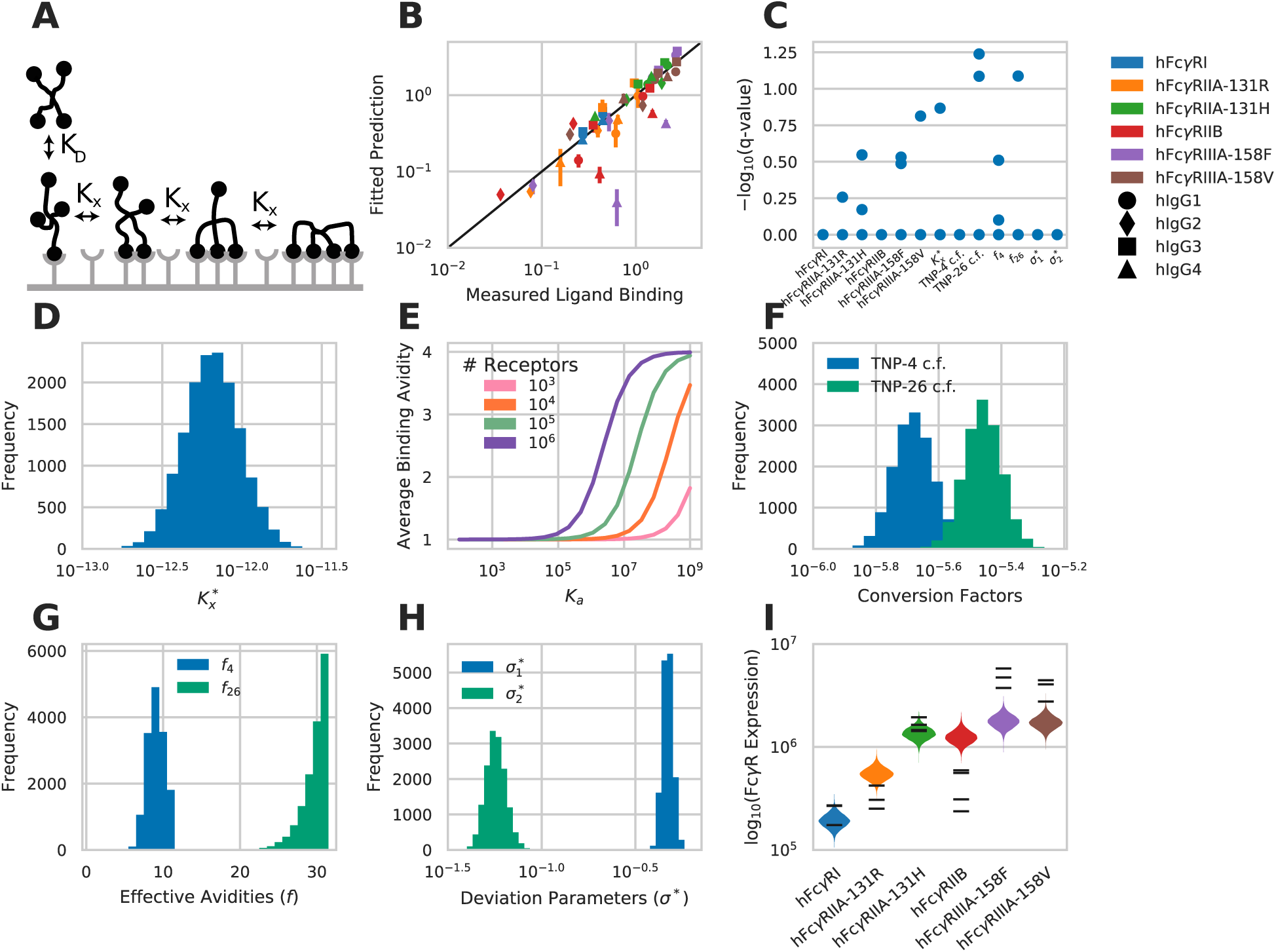
A multivalent binding model accounts for IgG-FcγR binding. A) Schematic of the multivalent binding model for interaction of an IC with a single species of hFcγR. B) Predicted versus measured binding for each hFcγR-hIgG pair at each avidity. C) Geweke convergence criterion for each walker of the MCMC chain. A significant p-value indicates failed convergence. D) Marginal distribution for the crosslinking constant *K*_*x*_^*∗*^. E) Average binding avidity predicted for a single interaction between a cell and an IC of avidity four, versus monovalent binding affinity at varied receptor expression levels. F) Marginal distribution for the constants to convert IC binding to normalized MFI. G) Marginal distribution for the avidities of TNP-4-BSA and TNP-26-BSA. H) Marginal distribution for each distribution spread parameter. I) The marginal distributions for receptor expression within each cell line expressing a single hFcγR subtype. Experimental measurements of receptor expression (Fig. 1B) are individually overlaid.

Treating *K*_*x*_ as constant across receptor-epitope combinthat there exists a crosslinking coefficientations may be reasonable when dealing with interactions of similar affinity, but clearly breaks down when a wide range of receptor-ligand affinities is involved. For example, for an FcγR-IgG interaction of barely measurable affinity, one would not expect to see multimeric binding occur with the same partitioning as an extremely high affinity interaction. Most concerning, given the assumption of equilibrium, is that an assumption of constant *K*_*x*_ violates detailed balance when this model is extended to include multiple receptor species that all bind the same epitope. Therefore, in attempt to resolve these issues, we assume that *K*_*x*_ is proportional to *K*_*a*_, such that there exists a crosslinking coefficient 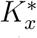 such that 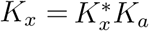 With this assumption, detailed balance is preserved, and *K_x_* is reduced when *K*_*a*_ is very low, as expected (see Methods).

We utilized a Markov chain Monte Carlo simulation to fit this model to our measurements of hFcγR-hIgG binding^19^. We observed close agreement between measured values and our model’s prediction of each condition (Fig. 2B). Both the sample autocorrelation (fig. S1) and the Geweke diagnostic (Fig. 2C) indicated sampling convergence^22^. Inspecting the fit of each parameter revealed that all parameters were well-specified, and that many of the parameter fits closely agreed with prior expectations. The 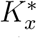 parameter was found to be roughly 10^−12^ (Fig. 2D). For the high-affinity hFcγRI-hIgG1 interaction, this comes out to a crosslinking constant of 6 x 10^−5^ per cell^−1^, close to the crosslinking constant of 1.35 x 10^−5^ per cell^−1^ observed for the high affinity human IgE-hFcɛR interaction^18^. This fit for the crosslinking coefficient falls within a range that provides differing binding behaviour dependent upon the affinity of the receptor interaction (Fig. 2E). Namely, the predicted binding valency upon a single IC interaction responds to changes in both affinity and receptor expression level within the range experimentally observed. Finally, there was a 1.6-fold difference in the fit values of the conversion coefficients that convert the number of TNP-4-BSA and TNP-26-BSA bound to mean fluorescent intensity, with the coefficient for TNP-26-BSA being larger (Fig. 2F). A difference in the IC-to-fluorescence conversion coefficients is expected as IC binding was measured with an anti-TNP fluorescent antibody.

Our prior distribution constrained the TNP-4-BSA and TNP-26-BSA avidities between one and twelve and between 20 and 32 respectively, which must be considered when interpreting the resulting fit (Fig. 2G). Within this constraint, there was a strong preference toward higher effective avidity with both species. The method for coupling TNP to BSA presumably creates a distribution of avidities and so deviation from the average is not surprising. A preference toward higher than average avidity binding is perhaps consistent with our earlier measurements that avidity has a potent effect on the level of binding (Fig. 1). That the deviation parameter for the IC binding data, 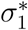 is fit to a greater value than the standard error of the receptor expression measurements, 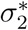 is consistent with greater variation in the former measurements (Fig. 2H). We compared 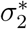 to its experimental value by calculating the standard errors of the receptor expression measurements normalized to the means of these measurements, then averaging these values. The resultant number (0.08) was close to and fell within one standard deviation of the predicted value (0.06). Finally, fits of receptor expression closely matched measured levels (Fig. 2I). In total, this demonstrates that a multivalent binding model accurately captures hFcγR-hIgG binding.

### A binding model provides specific predictions for the coordinate effects of IC abundance, avidity, and IgG subclass

With confidence that an equilibrium binding model can predict FcγR-IC binding, we sought to apply the model to make predictions about the combined influence of avidity and affinity on effector function. We focused on hFcγRIIIA, which is expressed alone within NK cells or alongside hFcγRIIB in dendritic and oter cell populations^10,23^. Predicted binding curves showed a shift left with increased avidity, consistent with the avidity-dependent shift observed experimentally elsewhere^5^ (Fig. 3A). Examining the number of receptor-receptor crosslinks and the abundance of receptors in multimer complexes showed a strong change with avidity, and biphasic concentration dependence (Fig. 3B-C). As a consequence of the differences in behavior between receptor multimerization and binding, increased avidity leads to far more oligomerization at a comparable level of receptor binding (Fig. 3B-D). For example, an IC of avidity 32 leads to the same amount of receptor oligomerization as the peak for avidity two at a 1000-fold lower concentration, and while binding roughly 50-fold less receptor. This emphasizes that avidity is an essential factor to consider for enacting effector responses and that measured receptor binding alone cannot predict consequent FcγR response.

**Figure 3:**
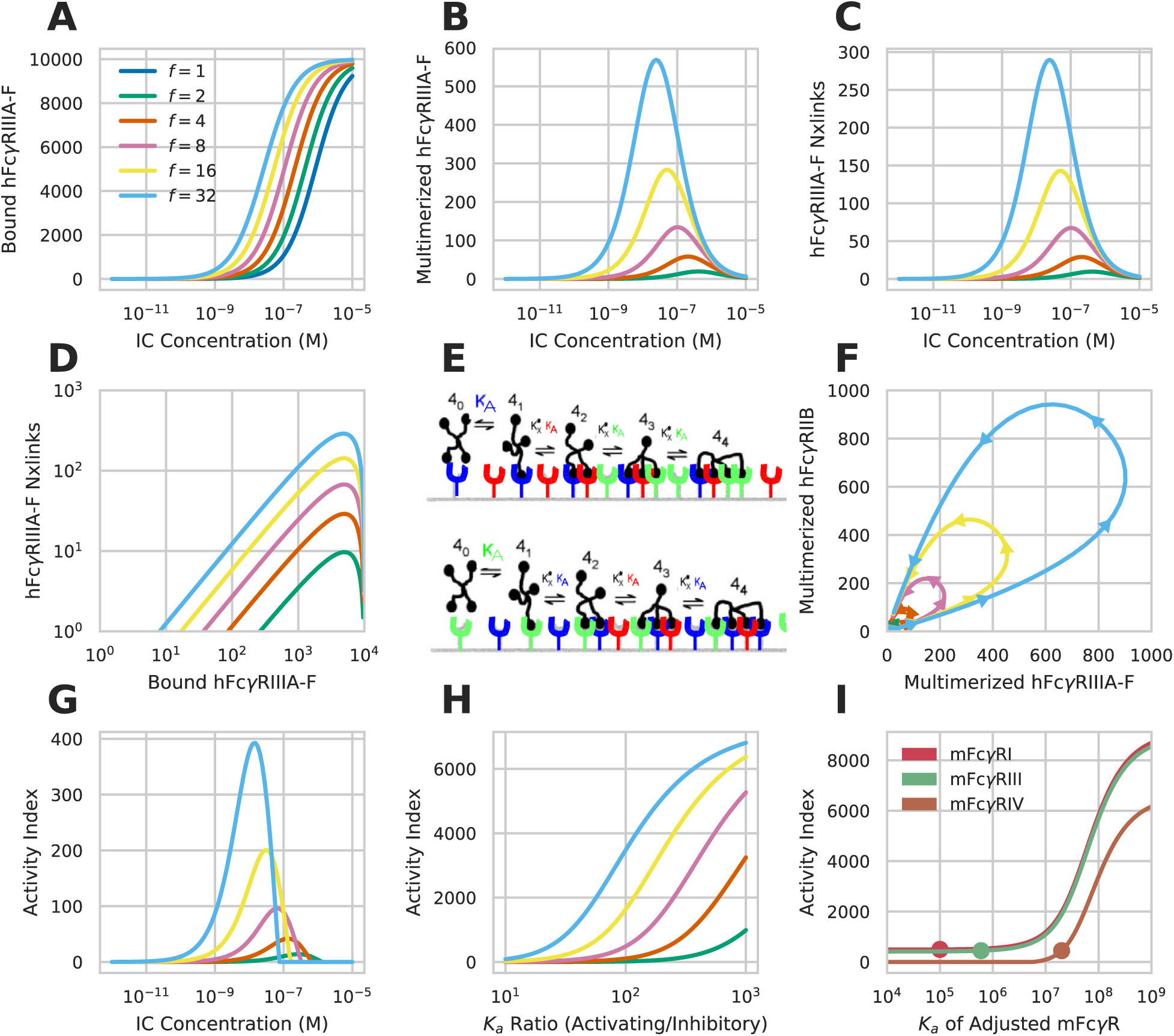
Specific predictions for the coordinate effects of IC parameters. A-C) Predicted hFcγRIIIA-F-hIgG1 binding (A), multimerized receptor (B), and number of receptor crosslinks (C) versus IC concentration at varied avidities. D) the amount of IC binding versus number of crosslinks for varied avidities. E) Schematic of the multivalent binding model for interaction of an IC with multiple species of FcγR. An individual IC can interact with a heterogeneous mix of receptors according to their affinities. The effective association constant for any crosslinking step, *K_x_*, is proportional to affinity. F) the predicted amount of multimerized receptor versus avidity for a cell expressing hFcγRIIIA-F and hFcγRIIB simultaneously when hIgG1-IC concentration is varied from 1 pM to 10 μM (beginning and ending near the origin). G) the calculated activity index (see Methods) for the conditions in F. H) Change in the activity index versus the A/I ratio for variations in hFcγRIIIA-F affinity responding to 1 nM hIgG1-ICs. I) Change in the activity index upon varying the affinity of mFcγRI, mFcγRIII, and mFcγRIV simultaneously expressed along with mFcγRIIB responding to 1 nM mIgG2b-ICs at an avidity of 5. Dot indicates the affinity of the receptor when not varied. Activity index increased by 50 at all values of *K_a_* for mFcγRI to make its curve visible.

In contrast to NK cells, other innate immune cell types express higher-affinity activating and lower-affinity inhibitory FcγRs in combination. Specifically, the inhibitory receptor mFcγRIIB is known to dampen effector function both endogenously and during a response to exogenous mIgG treatment^24^. We wondered how coordinate expression of an activating and inhibitory receptor would modulate the hFcγR-IC complexes formed. We first plotted the abundance of multimerized hFcγRIIIA versus multimerized hFcγRIIB for varied avidity and concentration of ICs. This indicated that, at low concentrations, hFcγRIIIA multimerization was higher at all avidities (Fig. 3F). Past a threshold concentration, hFcγRIIIA multimerization plateaued, and hFcγRIIB multimerization accumulated. This effect was observed for all avidities greater than one, only varying in the magnitude of the difference in multimerization, indicating that the effect of the lower affinity inhibitory receptor may modulate the concentration‐ and avidity-dependence only modestly. We additionally explored the influence of hFcγRIIB by defining an activity index (see Methods) as a surrogate measure for effector response. This also showed that inhibitory receptor has minor effects on the dose response relationship with respect to concentration or avidity (Fig. 3C vs. G).

Instead, we wondered if an inhibitory receptor plays a larger role on influencing the relative response between IgG subclasses as compared to IC concentration or avidity. Indeed, earlier work examining interventions of antibodies with constant variable regions but of differing IgG subclass identified that the ratio of the highest affinity activating receptor *K_a_* to that of the inhibitory receptor (A/I ratio) could predict the influence of each intervention. While this ratio has proven successful across many contexts, we wondered if our model might help identify the reason for this quantity’s ability to predict effector response^3,12,13^. By varying the affinity of the activating receptor, we indeed observed a strong relationship between the A/I ratio and activity index over the range of ratios previously examined (Fig. 3H). A second implicit assumption of the A/I ratio is that the highest affinity receptor is the primary activating receptor that influences response. By varying the affinity of each receptor in a four receptor model, we observed that the highest affinity receptor had the greatest influence on the activity index (see Methods) (Fig. 3I). Indeed, as lower affinity receptors were shifted to have higher affinity than the original highest affinity receptor, the response shifted toward dependence on that receptor.

### An IgG-FcγR binding model deconvolves *in vivo* function

We wished to explore whether a multivalent binding model can enable one to reverse engineer effector function *in vivo.* We posited that our modeling approach would allow one to predict the effect of therapeutic interventions involving ICs with defined IgG subclass composition on the effector responses of different cell populations based on their FcγR expression profile. Prior studies investigating treatments for HIV, cancer, and autoimmune dysfunction have utilized antibodies with identical variable regions, while varying other parameters of FcγR engagement, to elucidate the influence of these parameters on effector function^12,13^. One finding from these studies is that the relative affinity of an IgG subclass for each FcγR is important to the resulting response. We hypothesized that a more exact model of FcγR engagement would more robustly predict effector response (Fig. 4A).

**Figure 4:**
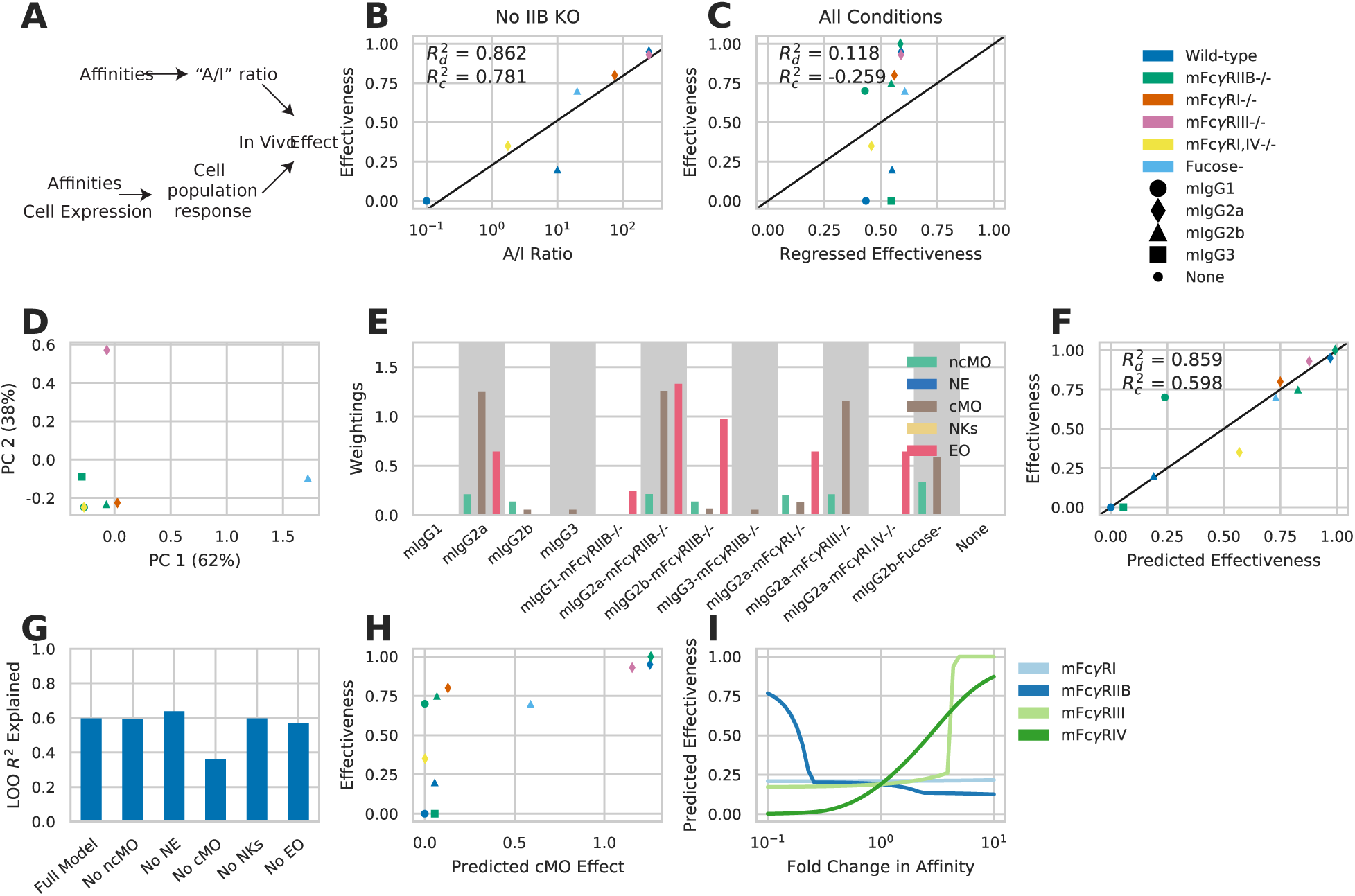
An FcγR-IgG binding model deconvolves *in vivo* function. A) Schematic of earlier IgG subclass experiments (top) and our approach (bottom). B) Effectiveness of individual mIgG interventions versus the A/I ratio for each mIgG constant region. Efectiveness is the percent reduction in lung metastases observed with treatment. C) Predicted versus regressed efectiveness for mIgG interventions upon mFcγR knockout using the maximal activating mFcγR affinity and inhibitory mFcγR affinity. D) Principal components analysis of the relevant affinities within each condition of mIgG treatment along with mFcγR knockout. Both axes scaled by a factor of 10^−8^. E) Individual quantities calculated for each intervention using receptor multimerization predicted by multivalent binding model and the activity index. Each quantity is scaled according to the weighting applied by the fted regression model. F) Efectiveness predicted by the multivalent binding model, quantifed by activity index, versus observed efectiveness. G) Leave-one-out model prediction R**^2^** with individual input components removed. H) Calculated activity index for cMO versus overall efectiveness of each intervention. I) Predicted effect of modulating each in-dvidual mFcγR affinity of mIgG2b. Predicted efectiveness increased by 0.02 for mFcγRI, at all values of fold change in affinity, to make its curve visible.

To study *in vivo* effector response, we focused on the manipulations made in one study wherein antibodies against the B16F10 melanoma antigen TRP1 (TA99) were applied to block lung metastasis in C57BL/6 mice^12^. Using a panel of antibodies with differing constant region but identical antigen binding revealed that an A/I ratio predicted response as observed previously (Fig. 4B). However, we noted that murine IgG2b (mIgG2b) showed divergence from the strong relationship observed with the other antibody constructs. To study this panel of interventions further, we included a number of murine FcγR (mFcγR) knockout or blocking manipulations by assuming the affinity of the receptor would be zero (SUPP TABLE). The A/I ratio cannot exist in the absence of an inhibitory receptor, and so we regressed the log-transformed maximal activating affinity and inhibitory receptor affinity, including knockout receptor conditions, to see whether this information could predict response in the broader panel (Fig. 4C). The highest affinity activating receptor *K*_*a*_and inhibitory receptor *K*_*a*_, even when treated as separate quantities, poorly predicted response in this larger panel, suggesting that other information is required to predict response within a wider panel of perturbations.

Examining the affinities present with each intervention emphasized that each antibody treatment differs in a multivariate way (Fig. 4D); a single principal component explained only 62% of the variation in affinities across each condition. We hypothesized that, although the A/I ratio captures the dominant variation for smaller changes in antibody binding, the other variation becomes important for more divergent interventions. Using mFcγR measurements for a panel of peripheral immune populations (fig. S2A), we applied our binding model to predict the activity index for each population and intervention, assuming a set avidity and ligand concentration (Fig. 4E). Regressing the activity index of each cell population against response showed greater predictive capacity than the A/I ratio within the wider panel of interventions (Fig. 4F). Multiple cell populations, including eosinophils and classical monocytes, were required for predicting response (Fig. 4G). Model prediction was robust to avidity and ligand concentration within an order of magnitude (fig. S2B).

The components of the resulting predictive model revealed how the A/I ratio is usually but not always predictive. Each input quantity was scaled by model weighting to indicate the relative contribution of each cell population (Fig. 4E). From this, the divergent response of mIgG2b was elucidated. mIgG2a is predicted to operate prominently through classical monocyte activity by our model (Fig. 4H), consistent with recent findings^25^. In contrast, mIgG2b and mIgG2b-fucose are predicted to operate through both types of monocytes. Finally, our model provides predictions for the efficacy of interventions, such as engineered variants of mIgG2b, with specifically altered mFcγR affinities (Fig. 4I).

## Discussion

A unified multivalent binding model accounting for IC avidity and affinity provides a framework for reasoning about how IC binding is affected by changing antibody-antigen binding or constant region composition (Fig. 2). Glycosylation forms and engineered mutants, in addition to variation in antigen binding, exponentially expand the repertoire of antibody variants possible. The quantitative model presented here provides a robust framework for reasoning about the contributions of each of these variables. Since it would not be feasible to experimentally explore all combinations of these variables, such a model is necessary for knowing which combinations would result in the most effective immunotherapeutic intervention. Further, a unified model for IC engagement may make it possible to consider the effects of antibody combinations in a rational manner.

In addition to enabling rationally designed immunotherapeutic interventions, our model of IC engagement makes it feasible to infer the factors contributing to the *in vivo* efficacy of existing immunotherapies (Fig. 4). In particular, we show through the application of our model to murine immunotherapy data that it can predict the effect of tumor-targeted antibodies better than an affinity ratio alone. This model additionally provides a number of specific, testable predictions: First, even modest reductions in inhibitory FcγR engagement may drastically increase the efficacy of mIgG2b, but only so long as activating receptor affinities are preserved (Fig. 4I). This ability to predict effector function for each individual cell population, given the kinetics of binding to each FcγR, will help to explore the complex changes constant region glyclosylation confers to antibody function^26^. Second, our model implicates cMO cells in the ability of TA99 mIgG2a to clear lung metastases. Given recent results showing that tumor location influences the relevant effector cell populations, this suggests that the optimal antibody design may be dependent upon tumor location^25^.

Applying this model can provide focus for future IgG engineering. For example, further development may inform application of antibody combinations of differing class for synergistic effector function. While we lack information about the effective avidity of TA99 engagement, more detailed characterization of its antibody binding might enable coordinate rational design of antibody class and antigen avidity. Our model is dependent upon FcγR abundance and the quantitative relationship between binding state and cell response; therefore, further refinement of where these receptors are expressed and how they sense IC engagement will improve our ability to study the *in vivo* environment. The versatility of antibody-based therapies ensure broad applicability of this approach to many diseases in which IgG effector function plays a key role, including the design of therapeutic antibodies for the treatment of infectious disease, autoimmune disorders, and other cancers.

More generally, these results demonstrate the ability of molecular models linked to data-driven inference to deconvolve *in vivo* function. Due to the baffling complexity of the immune system, model-driven design is vital to the advancement of rationally designed immunotherapies. Like Fc receptors, many innate immune receptors are extensively characterized in their interactions and knockout effects, and yet operate through combinatorial complexity due to diversity in protein species. As binding profiles and signaling mechanisms of these receptors become better quan-tifed, model-driven design will prove to be a powerful vehicle for further immune engineering advancement.

## Methods

All analysis was implemented in Python, and can be found at https://github.com/meyer-lab/FcgR-binding, release 1.0 (doi: 00.0000/arc0000000).

### Immune complex binding measurement

IC binding to hFcγRs was analyzed using Chinese hamster ovarian (CHO) cells stably expressing hFcγRs as previously described^10^. Receptor expression was quantified using antibodies against each hFcγR and flow cytometry measurement as previously described^10^. Briefly, ICs were generated by coincubation of 10 μg/ml anti-TNP hIgG and 5 μg/ml TNP-coupled BSA for 3 h with gentle shaking at room temperature. ICs were incubated with 100,000 CHO cells stably expressing hFcγRs for 1 h under gentle shaking at 4℃. Bound ICs were detected by flow cytometry using a PE-conjugated goat anti-human IgG F(ab’)_2_ fragment at 0.5 mg/ml (Jackson ImmunoResearch Laboratories). Each IC binding measurement was normalized to the average of all the points within that replicate. Receptor expression was quantified in terms of absolute number through comparison to fluorescence standards (QSC microspheres, Bangs Labs). Data were analyzed with Flow Cytometry Analysis Software (FlowJo) or FACSDiva Software.

### FcγR abundance measurement for primary cells

FcγR abundance was measured by flow cytometry from peripheral blood leukocytes of female C57Bl/6J mice under steady state conditions.

After erythrocyte lysis of anti-coagulated blood, cells were stained with antibodies to enable identification of cell types and for quantification of Fcγ receptors as listed in the Supplementary Information. Prior to addition of the staining antibodies cells were blocked with anti-FcγR antibodies to avoid unspecific binding to Fc receptors. Samples for quantification of FcγR4 were blocked with anti-FcγR2b/3 antibody clone 2.4G2. Samples for quantification of mFcγRI, mFcγRIIb or mFcγRIII were blocked with anti-FcγRIV antibody clone 9E9, since FcγRIV receptor has been shown to be a potential cause for unspecific binding of certain antibody isotypes in flow cytometry^27^.

A typical cell identification strategy was as follows: Cell aggregates were excluded by their forward light scatter (FSC) characteristics (area vs. height) and dead cells based on their capability for DAPI uptake. Leukocytes were identified by expression of common leukocyte marker CD45. Among those lymphocytes and myeloid cells were gated based on low side scatter (SSC) characteristics and absence of the Ly6G marker of neutrophilic granulocytes. NK cells were identified by low to intermediate CD11b expression together with expression of NK marker NK1.1. Among the highly CD11b-positive but NK1.1-negative cells, CD11b^high^ CD62L^high^ Gr-1^high^ classical and CD11b^high^ CD62L^low^ Gr-1^low^ non-classical monocytes were distinguished based on their differential expression of Gr-1 and in most experiments additionally also of CD62L. Among the granulocytes with high SSC and CD11b expression, eosinophils were identified by their very high SSC, low FSC and absence of Ly6G, whereas neutrophils were characterized by intermediately high side and forward scatter and presence of the neutrophil marker Ly6G.

Surface receptor was quantitated by measuring antibody binding capacity (ABC) for antibodies specific for the respective Fcγ receptor. Calculation of ABC on cells is based on a reference curve for the correlation between fluorescence intensity (caused by the respective anti-FcγR antibody) and the number of antibody binding sites using a group of beads with known ABC values, reflecting their ability to bind a known amount of antibody. These curves were established in each experiment for all tested anti-FcγR antibodies, using Quantum Simply Cellular (QSC) anti-mouse or anti-rat beads (Bangs Laboratories Ltd.)—depending on the host species of the respective anti-FcγR antibody—according to manufacturer’s instructions. Anti-FcγR4 antibody 9E9 is derived from Armenian hamster, but was found to be efficiently bound by the anti-mouse QSC beads. Yet, in most experiments these beads were pre-coated with mouse anti-hamster moieties prior to staining with 9E9.

All anti-FcγR antibodies used for Fc quantification were used as conjugates with R-PE. They were either purchased pre-labelled or were conjugated in-house. In each experiment QSC microspheres are stained with the respective anti-FcγR antibody in the same concentration as it was used for cell staining. To allow subtraction of ABC-background accounting for background fluorescence of the cells, FMO (“fluorescence-minus-one”) controls were used in each experiment, where cells were stained with all antibodies for cell type identification but without anti-FcγR antibody. Flow cytometric analysis was done on a FACS Canto II (BD Biosciences, Heidelberg) and data were analyzed with FACSDiva Software (BD). For ABC calculation we used the QuickCal Software provided by Bangs Laboratories. FcγRs with average abundances less than 10^3^ per cell were considered absent; in every case this determination was consistent with measurements in samples from knockout animals.

### *In vivo* regression

Regression against *in vivo* effectiveness of mIgG treatments was performed by least-squares (scipy.optimize.least_squares). Association constants for all combinations of mIgG and mFcγR were obtained from previous experimental measurements^8,23^. Each effectiveness was presented as the percent reduction in the number of lung metastases quantified^12^. To account for the limited range of this quantity (e.g. one cannot have a reduction of 200%), the regression was transformed by tanh such that:

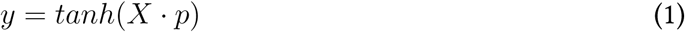

so that lim_x→∞_ *y*(*x*) = 1. *X* is the predicted mFcγR activity for each cell line according to our model, and *p* is the regression weights.

### PCA of mlgG-mFcγR affinities

Principle component analysis was performed using scikit-learn and the affinities of the four mIgGs (mIgG1, mIgG2a, mIgG2b, mIgG3), with or without knockout treatments, for each of four receptors (mFc γ RI, mFcγRIIB, mFcγRIII, mFcγRIV). Association constants for all combinations of mIgG and mFcR were obtained from previous experimental measurements^7^. The affinity for a knocked-out or blocked mFcγR was assumed to be zero. The affinities were not normalized before PCA transformation.

## Model

### Base model

TNP-BSA equilibrium binding to FcγRs was modeled using a two-parameter equilibrium model of multivalent ligand binding to monovalent receptors expressed uniformly on a cell surface^14,17^. This model assumes that the IC effectively presents a single kind of epitope and that the cell expresses exactly one receptor species that recognizes the epitope. The model also assumes an excess of ligand, such ligand concentration is effectively constant. Within the model, the initial binding of an IC to the cell is assumed to occur according to a monovalent binding interaction governed by the individual binding site association constant *K*_*a*_. Once a ligand is bound to the cell surface by one receptor, all subsequent binding occurs through crosslinking events with equilibrium partitioning *K*_*x*_, in which an unbound epitope on the ligand binds to a free receptor on the cell surface. *K*_*x*_ serves as the association constant for all crosslinking interactions. According to the model, the number of ligand bound *i*-valently to the cell at equilibrium, *v*_*im*_, can be found using the relation

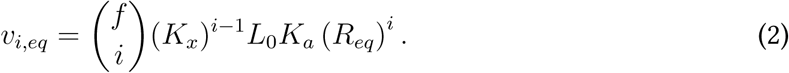

Here, *f* is the effective avidity of the ligand, *K*_*x*_ is a crosslinking parameter with units of # per cell, *L*_0_ is the concentration of ligand, and *R*_*eq*_ is the number of unbound receptors at equilibrium. Consequently, the total number of ligand bound at equilibrium is

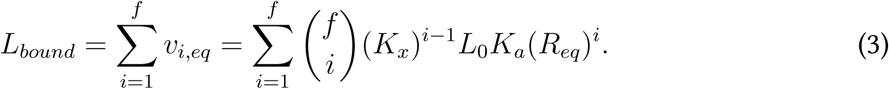

*R*_*eq*_ changes as a function of *f*, *L*_0_, *K*_*a*_, *K*_*x*_, and *R*_*tot*_, the total number of receptors expressed on the cell surface. It can be solved for numerically using the relationship

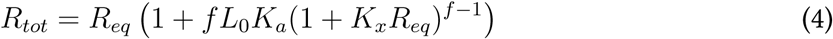

when these parameters are known. As a consequence of eq. 2, the number of receptors that are clusthered with at least one other receptor at equilibrium (*R*_*multi*_) is equal to

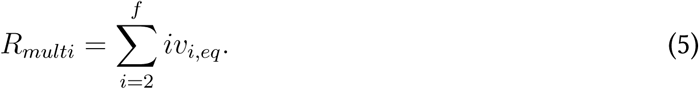

### Specification for *K*_*x*_

We represented *K*_*x*_ for any given crosslinking interaction as the product of *K*_*a*_, the affinity of the epitope being bound for the receptor species to which it binds, and a crosslinking coefficient, 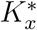, that is uniform for all combinations of FcγR and IgG. For any given crosslinking interaction between an epitope-receptor pair with affinity *K*_*a*_,

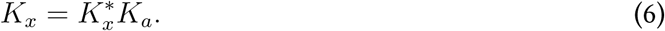

As a consequence of this construction, *K*_*x*_ becomes zero in the absence of binding and satisfies detailed balance.

### Parameters and assumptions

Association constants for all combinations of hIgG and hFcγR were obtained from previous experimental measurements^7^. In each replicate of the binding assay, cells were coincubated with 5 μg/ml TNP-4-BSA or TNP-26-BSA. Because the molar masses of 2,4,6-trinitrophenyl groups and of BSA are approximately 173 Da and 66 kDa, respectively, we represented the molar concentrations of TNP-4-BSA and TNP-26-BSA as 74 nM and 70 nM^10^. We also assumed that there were two different conversion factors for TNP-4-BSA and TNP-26-BSA between the number of ICs bound and the MFIs measured in the assay, due to IC detection occuring through TNP quanti-tation. Lastly, we assumed that, due to steric effects, the effective avidities of TNP-4-BSA and (especially) TNP-26-BSA might be different than their actual avidities. This required us to fit the following eleven parameters: the total expression level *R_tot_* for each hFcγR, 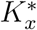, conversion factors from ligand bound to MFI measured for both TNP-BSAs, and effective avidities for both TNP-BSAs (*f*_4_ and *f*_26_, respectively). In our simulation, receptor expression levels were allowed to vary between 10^3^ and 10^8^, *K*_*x*_^*∗*^ between 10^−25^ and 10^3^ (in order to provide no constraint on possible values), the conversion factors between 10^−10^ and 10^5^, *f*_4_ between one and twelve, and *f*_26_ between twenty and thirty-two.

### Model ftting and deviation parameters

We fit a standard deviation parameter 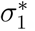. The likelihood for each combination of predicted values for 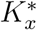 the two conversion factors, *f*_*4*_, and *f*_*26*_ for each hFcγR-hIgG-TNP-BSA combination was calculated by comparison of our experimental data to a normal distribution with mean equal to our model’s predicted binding and standard deviation equal to the predicted binding times 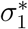.

In addition to IC binding, the receptor expression of each cell line was quantitatively measured. We assumed that the receptor expression measurements were log-normally distributed, with the standard deviation of the log-normal distribution being proportional to the common logarithm of the actual expression. We fit a second standard deviation parameter, 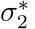 such that the likelihood of each receptor measurement was calculated using a normal distribution with mean equal to the common logarithm of the predicted receptor expression and standard deviation equal to this value times 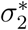. The overall likelihood of the model at each parameter set was calculated as the product of all individual likelihoods. 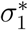 and 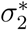 were allowed to vary between 10^−4^ and 10. The prior for each fitted parameter was therefore specified to be uniform on a log scale within its lower and upper bounds.

We fit our model to binding measurements for each hFcγR-hIgG pair using an affine invariant Markov chain Monte Carlo sampler as implemented within the emcee package^19^. We assayed the convergence of the Markov chain using the Geweke diagnostic and chain autocorrelation^22^. The Geweke diagnostic was used to determine whether early and late segments of the Markov chain could have been sampled from the same probability distribution. Each walker’s series of values for a particular parameter was treated as a single chain, upon which the diagnostic was evaluated.

### Generalized Multi-Receptor Model

To account for cells expressing multiple FcγRs, we extended the model to account for binding in the presence of multiple receptors. At each crosslinking step, *K*_*x*_ must be proportional to the *K*_*a*_ of the corresponding monovalent epitope-receptor interaction to satisfy detailed balance. For any cell expressing *N* distinct receptor species that all bind the same epitope, let *R*_*tot,i*_ be the total number of receptors *i* expressed on the cell surface, and let *K*_*a,i*_ be the affinity of receptor *i* for the epitope. Let IC, ligand, and 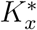 be as previously described (see Base Model). For all *i* in {1, 2,…, *N*}, let

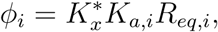

where *R*_*eq,i*_ is the number of receptors *i* unbound at equilibrium. The individual IC-receptor interactions of an IC, with effective avidity *f*, bound to *_qi_* receptors *i*, _*qj*_ receptors *j,* etc. can be represented by the vector **q** = (*q*_*1*_ *q*_*2*_,…, q_N+1_), where *q*_*N+1*_ is equal to the number of unbound epitope on the IC. For any such vector describing the binding state of an IC-receptor complex, the number of ICs bound in such a way at equilibrium is equal to

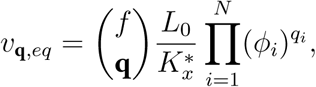

where 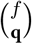 represents the multinomial coefficient 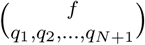. Therefore, for all receptors *i*, we have that *R*_*eq,i*_ satisfies the relation

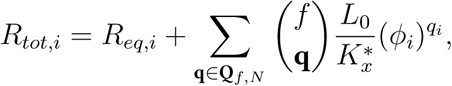

Where

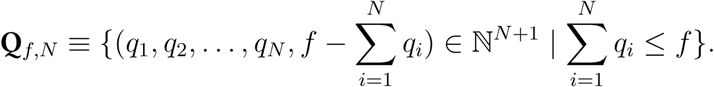

For our analysis, *R*_*eq,i*_ was solved for for all *i* by iterative root-finding using this relation, utilizing the Brent routine (scipy.optimize.brenth). Consequent of, the total number of ligand bound at equilibrium is

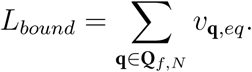

The number of receptor *i* that are multimerized at equilibrium can be calculated as

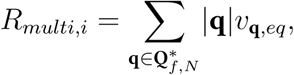

where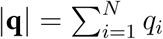and

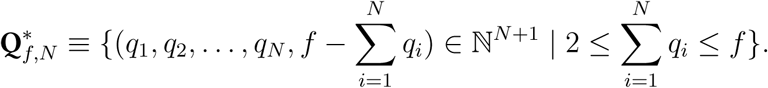

### Activity Index

To account for the combined effects of activating, inhibitory, and decoy receptors, we defined an activity index. To do so, we defined the activity index as being the dot product of the vector *v*, or number of multimerized receptors of each receptor species, and *w*, the activity of each receptor species. Activating receptors were given an activity of 1, decoy receptors 0, and inhibitory receptors −1. Multimerization states that resulted in activities of less than 0 were set to 0. This definition satisfied our expectations that activity increases with a greater number of activating receptors, decreases with more inhibitory receptors, and does not change with variation in the number of decoy receptors.

## Acknowledgements

Tis work was supported by NIH DP5-OD019815 to A.S.M., a Terri Brodeur Breast Cancer Foundation Fellowship to A.S.M., DFIG-CRC1181-A07 to F.N., DFIG-TRR130-P13 to F.N., and in part by the Koch Institute Support (core) grant P30-CA14051 from the NCI. The authors wish to thank Song Yi Bae, Simin Manole, and Ted Richards for helpful feedback. **Competing fnancial interests:** The authors declare no competing financial interests.

## Author contributions statement

A.S.M. and F.N. conceived the experiment(s), A.S.M., R.A.R., M.B., N.G., and A.L. conducted the experiments and analyzed the results. All authors reviewed the manuscript.

## Supplement

**Murine knockout conditions and mFcγR-mIgG affinities.** Murine knockout conditions and mFcγR-mIgG affinities used in principal component analysis shown in Fig. 4. Values in columns labeled with mFcγRs are affinities with M^−1^ units^89^. Values in the effectiveness column represent proportional reduction in lung metastases observed with treatment^12^.

| Condition | mFcγRI | mFcγRIIB | mFcγRIII | mFcγRIV | Effectiveness |
| --- | --- | --- | --- | --- | --- |
| mIgG1 | 0.0 | $3.0 \times 10^6$ | $3.0 \times 10^5$ | 0.0 | 0.00 |
| mIgG2a | $1.0 \times 10^8$ | $4.0 \times 10^5$ | $7.0 \times 10^5$ | $3.0 \times 10^7$ | 0.95 |
| mIgG2b | $1.0 \times 10^5$ | $2.0 \times 10^6$ | $6.0 \times 10^5$ | $2.0 \times 10^7$ | 0.20 |
| mIgG3 | $2.0 \times 10^7$ | 0.0 | 0.0 | 0.0 | 0.00 |
| mIgG1-FcγRIIB-/- | 0.0 | 0.0 | $3.0 \times 10^5$ | 0.0 | 0.70 |
| mIgG2a-FcγRIIB-/- | $1.0 \times 10^8$ | 0.0 | $7.0 \times 10^5$ | $3.0 \times 10^7$ | 1.00 |
| mIgG2b-FcγRIIB-/- | $1.0 \times 10^5$ | 0.0 | $6.0 \times 10^5$ | $2.0 \times 10^7$ | 0.75 |
| mIgG3-FcγRIIB-/- | $2.0 \times 10^7$ | 0.0 | 0.0 | 0.0 | 0.00 |
| mIgG2a-FcγRI-/- | 0.0 | $4.0 \times 10^5$ | $7.0 \times 10^5$ | $3.0 \times 10^7$ | 0.80 |
| mIgG2a-FcγRIII-/- | $1.0 \times 10^8$ | $4.0 \times 10^5$ | 0.0 | $3.0 \times 10^7$ | 0.93 |
| mIgG2a-FcγRI,IV-/- | 0.0 | $4.0 \times 10^5$ | $7.0 \times 10^5$ | 0.0 | 0.35 |
| mIgG2b-Fucose-/- | $1.0 \times 10^5$ | $1.0 \times 10^7$ | $1.1 \times 10^6$ | $2.0 \times 10^8$ | 0.70 |

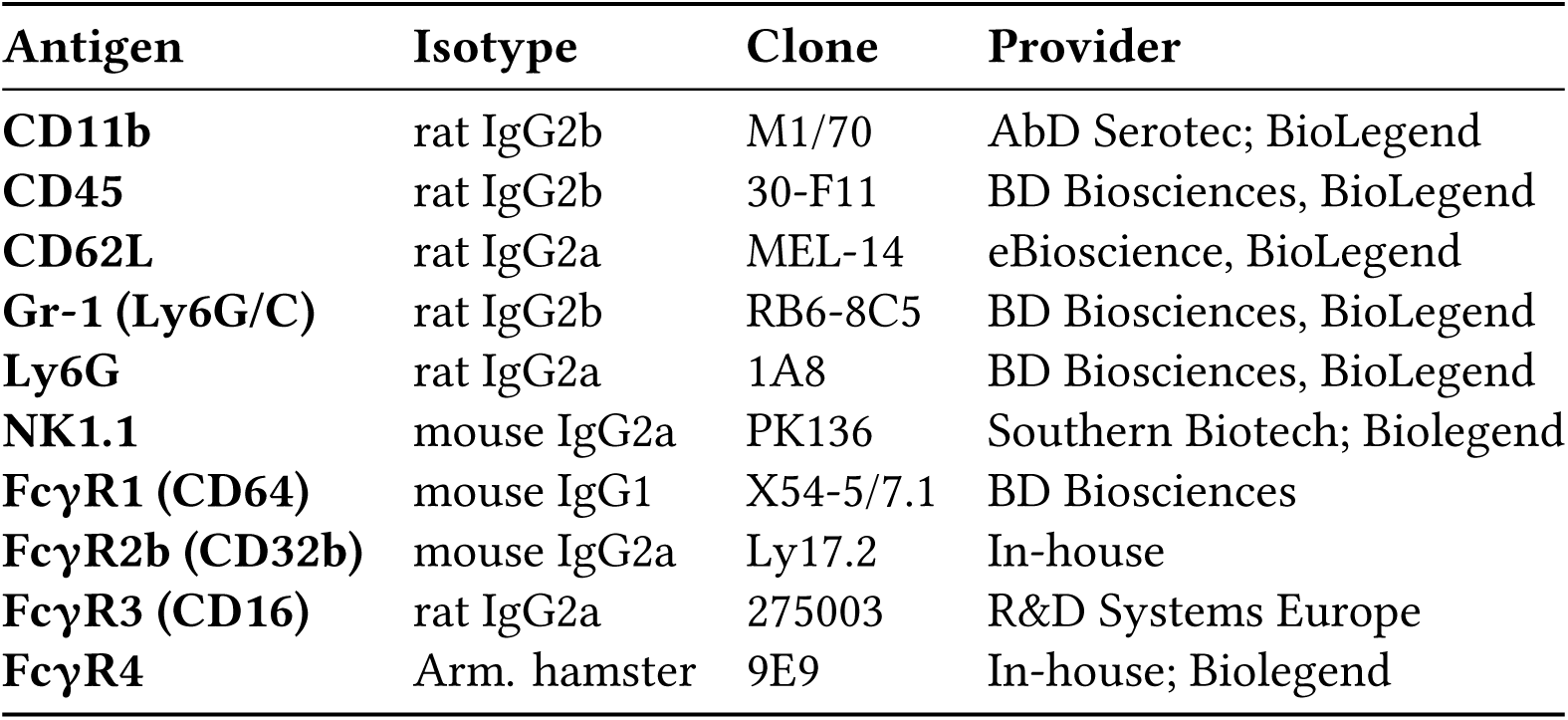
Antibodies for cell identification and mFcγR quantitation.

| Antigen | Isotype | Clone | Provider |
| --- | --- | --- | --- |
| **CD11b** | rat IgG2b | M1/70 | AbD Serotec; BioLegend |
| **CD45** | rat IgG2b | 30-F11 | BD Biosciences, BioLegend |
| **CD62L** | rat IgG2a | MEL-14 | eBioscience, BioLegend |
| **Gr-1 (Ly6G/C)** | rat IgG2b | RB6-8C5 | BD Biosciences, BioLegend |
| **Ly6G** | rat IgG2a | 1A8 | BD Biosciences, BioLegend |
| **NK1.1** | mouse IgG2a | PK136 | Southern Biotech; Biolegend |
| **FcγR1 (CD64)** | mouse IgG1 | X54-5/7.1 | BD Biosciences |
| **FcγR2b (CD32b)** | mouse IgG2a | Ly17.2 | In-house |
| **FcγR3 (CD16)** | rat IgG2a | 275003 | R&D Systems Europe |
| **FcγR4** | Arm. hamster | 9E9 | In-house; Biolegend |

**Figure S1:**
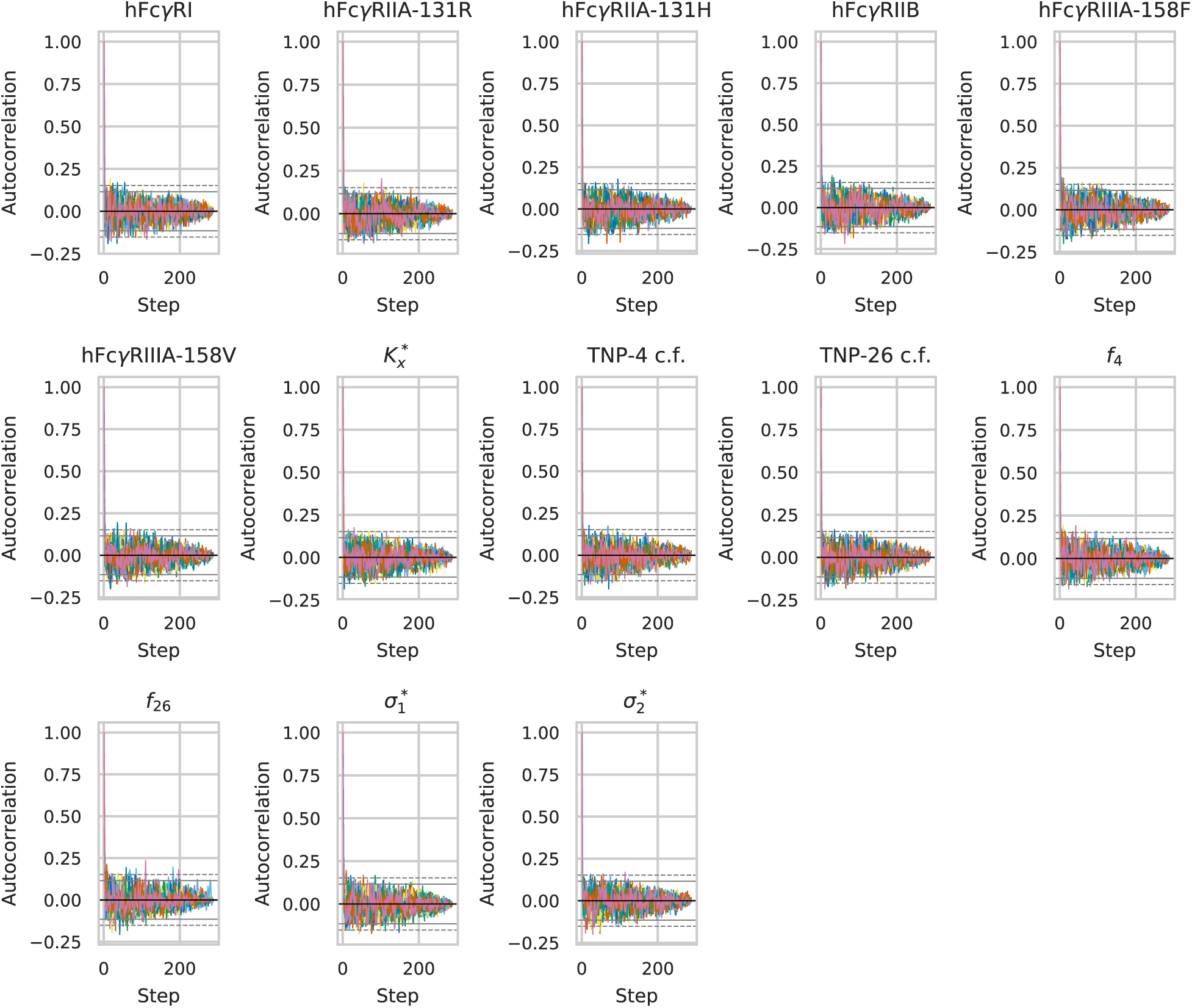
Parameter autocorrelation for each parameter and each walker. Solid and dotted lines indicate the 95th and 99th percentile significance bounds

**Figure S2:**
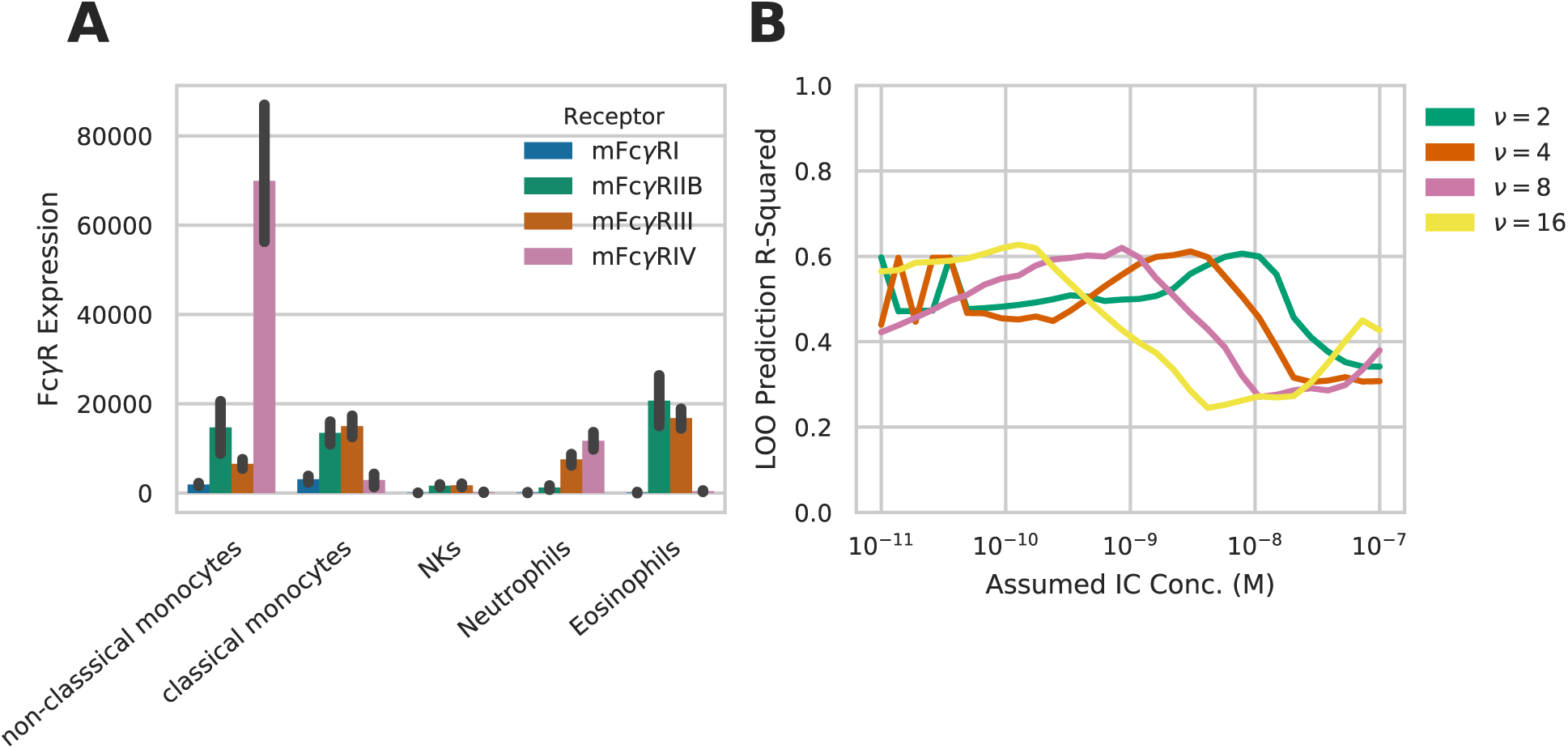
Supplementary results to *in vivo* predictions. A) Receptor abundance quantified across different peripheral immune cell populations. Error bars indicate standard error of biological replicates (N >= 3).B) Prediction performance upon crossvalidation for the model predicting *in vivo* efficacy with differing assumed IC concentration and avidity

